# An Analysis of SARS-CoV-2 Using ViReport

**DOI:** 10.1101/2020.06.20.163162

**Authors:** Miranda Song, Niema Moshiri

## Abstract

The ongoing outbreak of *severe acute respiratory syndrome coronavirus 2* (SARS-CoV-2) has resulted in millions of cases and hundreds of thousands of deaths. Given the current lack of treatments or vaccines available, it may be useful to trace the evolution and spread of the virus to better develop methods of preventative intervention. In this study, we analyzed over 4,000 full genome sequences of human SARS-CoV-2 using novel tool ViReport [13], an automated workflow for performing phylogenetic analyses on viral sequences and generating comprehensive molecular epidemiological reports. The complete ViReport output can be found at https://github.com/mirandajsong/ViReport-SARS-CoV-2.

## 1 Introduction

Coronaviruses are a group of RNA viruses that cause various diseases in humans, other mammals, and birds. In the last two decades, there have been three documented strains of coronavirus that have caused major epidemics in humans. These include *severe acute respiratory syndrome coronavirus* (SARS-CoV) in 2002, *Middle East respiratory syndrome coronavirus* (MERS-CoV) in 2012, and most recently, a novel coronavirus referred to as COVID-19 or SARS-CoV-2 that emerged in Wuhan, China in December 2019. As of June 2020, there have been over 8 million confirmed cases of SARS-CoV-2 globally [3].

The rapid spread of SARS-CoV-2 has demonstrated the need to understand how the virus spreads and for measures of epidemic intervention such as stay-at-home orders and social distancing. To gain insight into disease transmission and origins, phylogenetic analyses are often performed using bioinformatics methods to quantify and visualize the relatedness of viral sequences. A typical workflow for phylogenetic analysis and reconstruction involves four main steps: (1) Data Acquisition, (2) Sequence Alignment, (3) Phylogenetic Inference, and (4) Tree Visualization [6].

Acquiring data involves identifying homologous nucleotide or protein sequences that are representative of the point of interest, whether that be a specific gene or species or something else. This step may involve additional preprocessing (e.g. filtering by length or sample type) to ensure that the sequences are valid and that the inferred tree will provide meaningful results. There are many public databases that provide viral (and other) sequences and accompanying metadata, including NCBI (ncbi.nlm.nih.gov), GISAID (gisaid.org), and ViPR (viprbrc.org).

Multiple sequence alignment (MSA) refers to the alignment of related biological sequences of similar length, the output of which may be used to infer more specific evolutionary relationships. Given the classification of MSA computation as an NP-complete problem, there are many existing algorithms that solve this problem [1]. Examples include MAFFT [7], MUSCLE [5], and ViralMSA [15], all of which are implemented in ViReport, along with others.

Phylogenetic inference is the reconstruction of evolutionary trees by grouping individual elements based on shared ancestry. Estimation of phylogenetic trees can be performed through various methods such as Bayesian Inference and Maximum Likelihood [6], which have spawned tools such as BEAST [4] and RAxML [17], respectively. The resulting phylogenies can be input into online tree visualization tools for further examination.

While there exist various tools that seek to automate this process such as MEGA5 [18] and Phylogeny.fr [2], ViReport is robust in its integration of modules that allows for the user to select their desired tool for each stage in the workflow. Furthermore, ViReport includes options for rooting and dating the inferred phylogeny, calculating pairwise distances, ancestral sequence reconstruction, and transmission clustering, all of which provide additional information beyond the tree itself.

**Figure 1:**
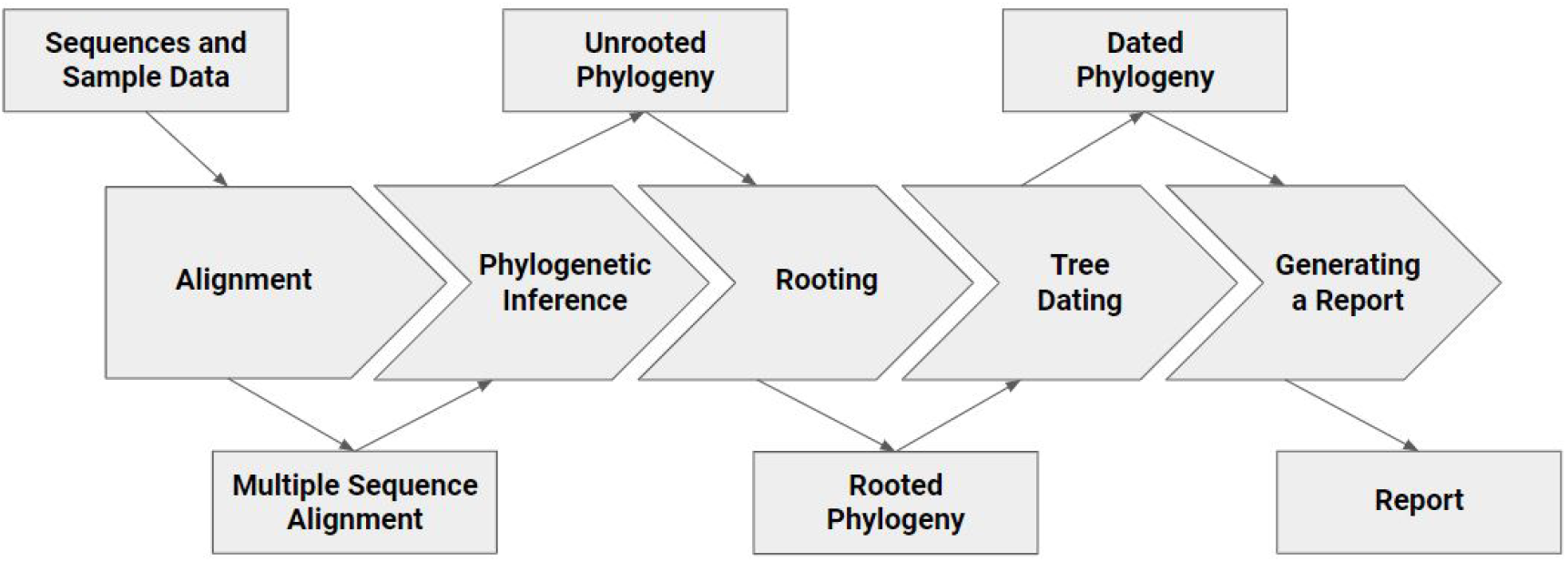
Demonstrates the overall ViReport workflow.

## 2 Methods

ViReport was used to preprocess the data, perform the MSA and phylogenetic inference steps, and produce the input tree files for tree visualization.

### 2.1 Data Acquisition

For this study, sequences were retrieved from the NCBI Virus database, which hosts data from GenBank. Nucleotide sequences were selected for using the SARS-CoV-2 Virus specification and by filtering out sequences under 26,000 bases as to only include full genome sequences [22]. Sequence metadata was also compiled, specifically that regarding the coun-try of origin and the collection date.

### 2.2 Multiple Sequence Alignment

Multiple Sequence Alignment was performed using ViralMSA [15] utilizing Minimap2 [10] and the reference sequence with accession ID NC 045512, defined as “Severe acute respiratory syndrome coronavirus 2 isolate Wuhan-Hu-1” and dated December 2019. Pair-wise sequence distances were calculated using the tn93 tool from HIV-TRACE [8].

### 2.3 Phylogenetic Inference

FastTree 2 [16] was used to infer a maximum-likelihood phylogeny under the General Time-Reversible model [19] using a Gamma20-based likelihood. The phylogeny was MinVar-rooted using FastRoot [11] and dated using LSD2 [20]. Pairwise phylogenetic distances were calculated using TreeSwift [14].

### 2.4 Tree Visualization

The phylogenies were visualized using iTOL [9] and IcyTree [21].

## 3 Results

### 3.1 Analysis of Different Coronavirus Strains

As a preliminary analysis based on a previous study titled *A Novel Coronavirus from Patients with Pneumonia in China, 2019* [23], 30 full genome sequences of different coronavirus strains (including SARS, MERS, and SARS-CoV-2) were compared using ViReport. After preprocessing, the average sequence length was 30018.133 bases with standard deviation 584.016, with sample dates ranging from April 2003 to March 2020. The average pairwise sequence distance (as calculated using tn93 from HIV-TRACE) was 0.503, with a standard deviation of 0.322.

**Figure 2:**
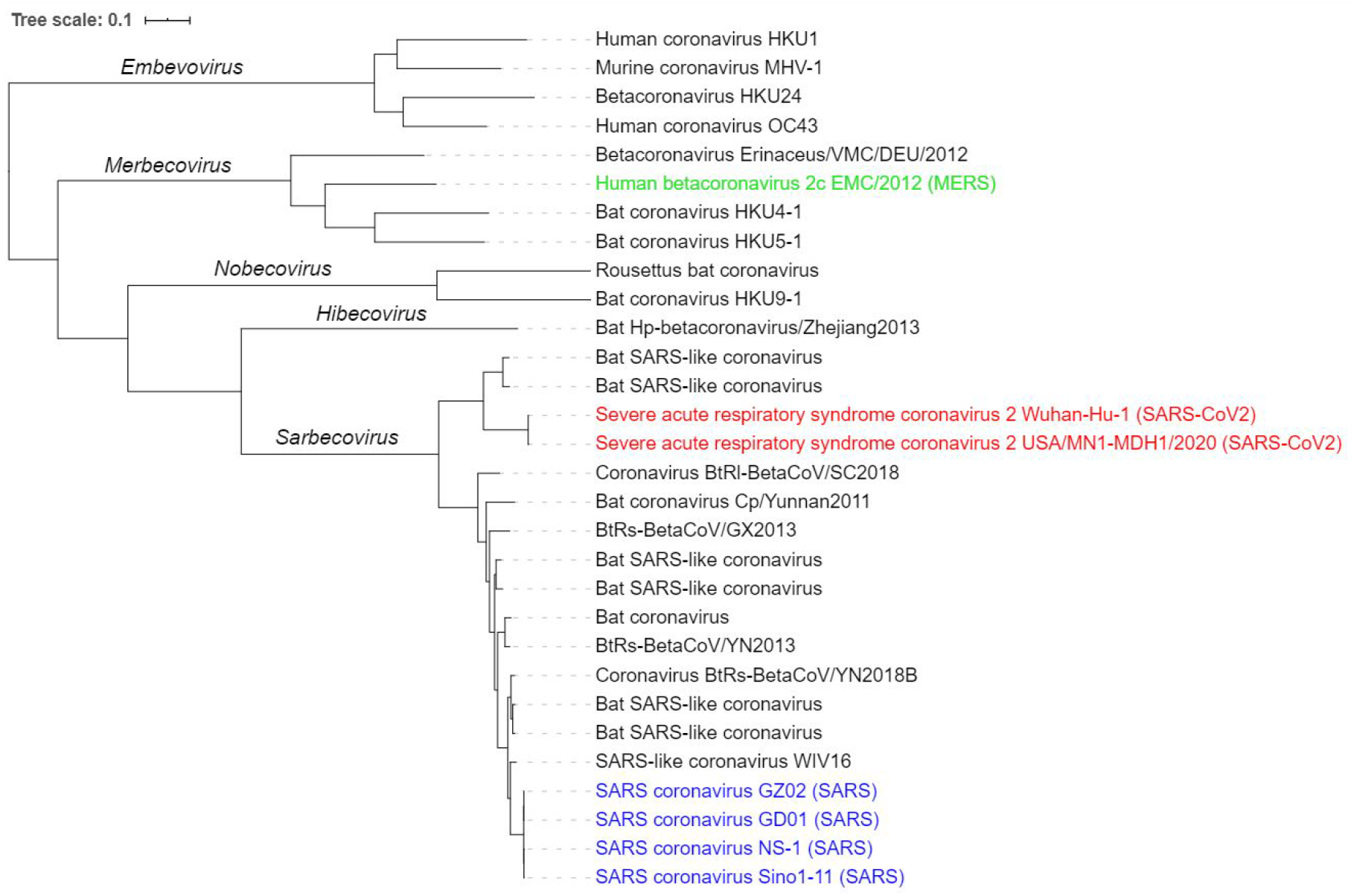
Rooted phylogenetic tree showing evolutionary relationships between various human and nonhuman coronavirus strains as produced by ViReport and visualized using iTOL; highlighted are SARS genomes in blue, MERS in green, and SARS-CoV-2 in red.

**Figure 3:**
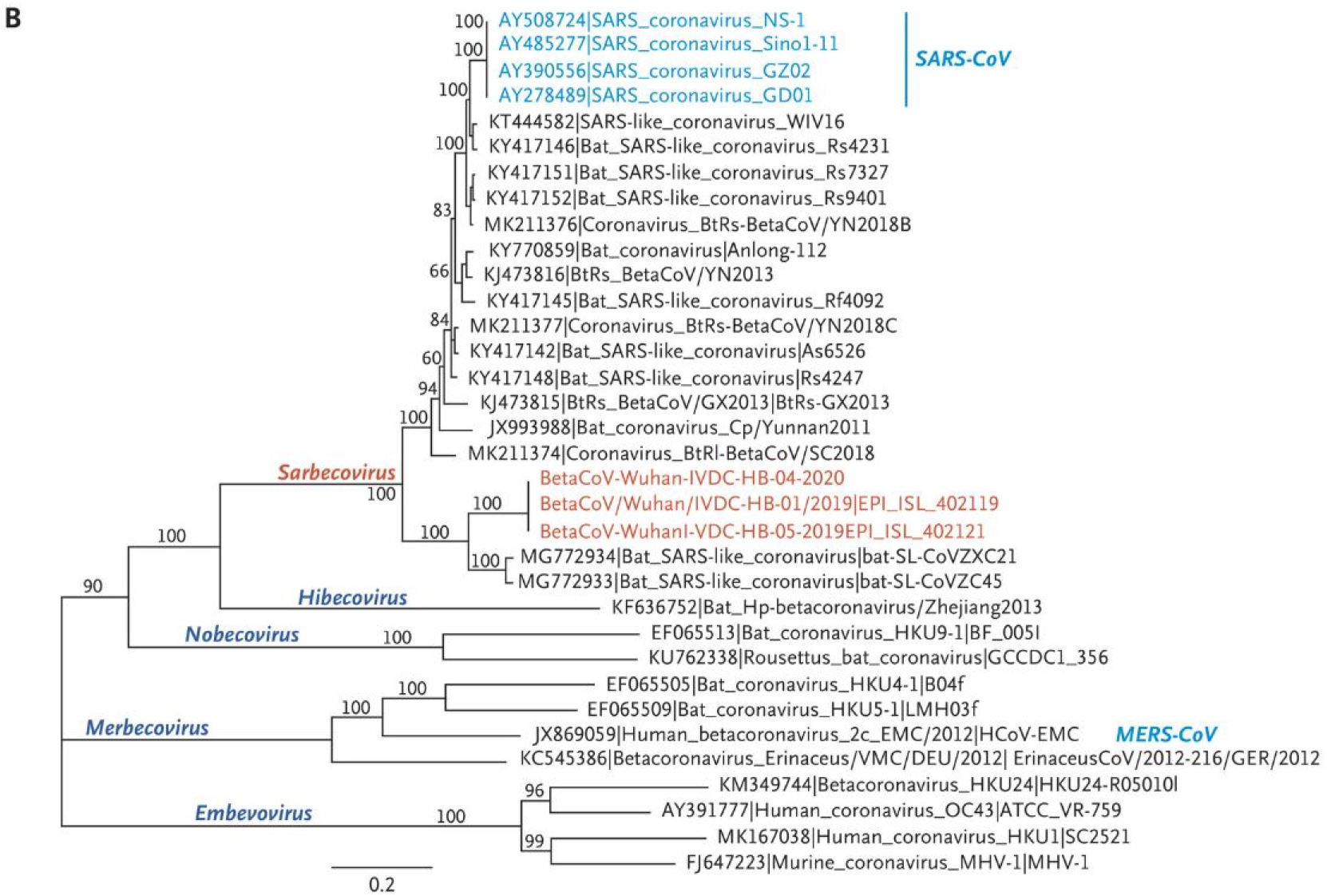
A reproduction of Figure 4B, “Phylogenetic Analysis of 2019-nCoV and Other Betacoronavirus Genomes,” from *A Novel Coronavirus from Patients with Pneumonia in China, 2019* [23]

Comparison of the visualization of the ViReport output and the figure from this study [23] shows that the results are very similar, despite the two analyses having used different tools (ViralMSA and IQ-TREE for ViReport, MUSCLE and RAxML for the article [23]), as the overall clades are nearly identical. Both analyses indicate that, while SARS-CoV, MERS, and SARS-CoV-2 fall into distinct clades, SARS-CoV-2 is more closely related to SARS-CoV, as both fall under the subgenus *Sarbecovirus*, whereas MERS falls under a different subgenus *Merbecovirus*. Additionally, SARS-CoV-2 is most closely related to a SARS-like coronavirus found in bats.

### 3.2 Analysis of SARS-CoV-2 Genomes

This analysis was performed on a dataset of 4635 full genomes sequences of SARS-CoV-2 found in *Homo sapiens*. The average sequence length was 29819.915 bases with a standard deviation of 80.154, with sample dates ranging from December 2019 to May 2020 (sequence data was retrieved on May 30, 2020; additional sequences have been added to the database since).

**Figure 4:**
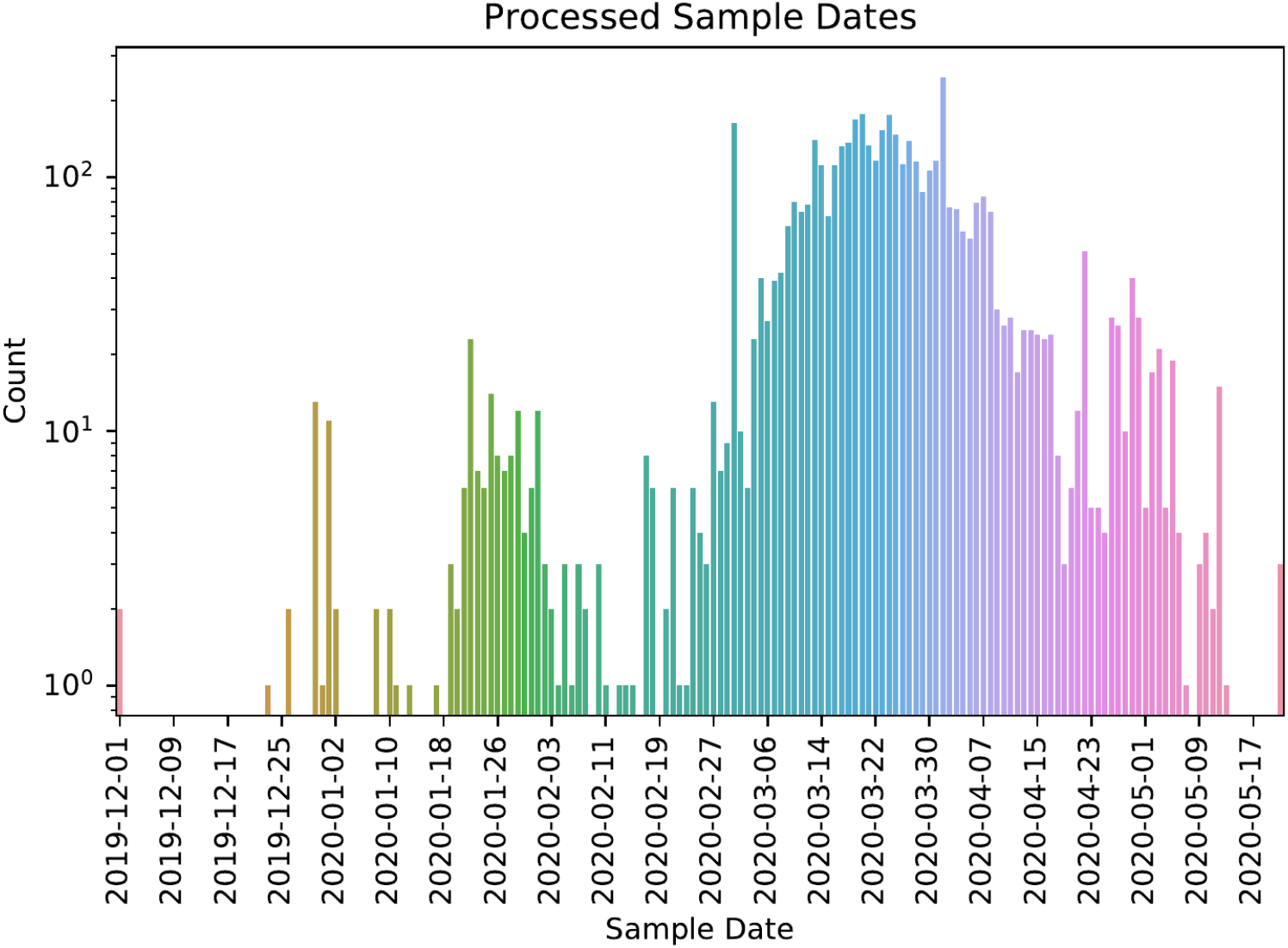
Distribution of sample dates.

We can see from the histogram above that the peak of the sample distribution falls around late March 2020. Sequences were also grouped by geographical location, as shown below.

**Figure 5:**
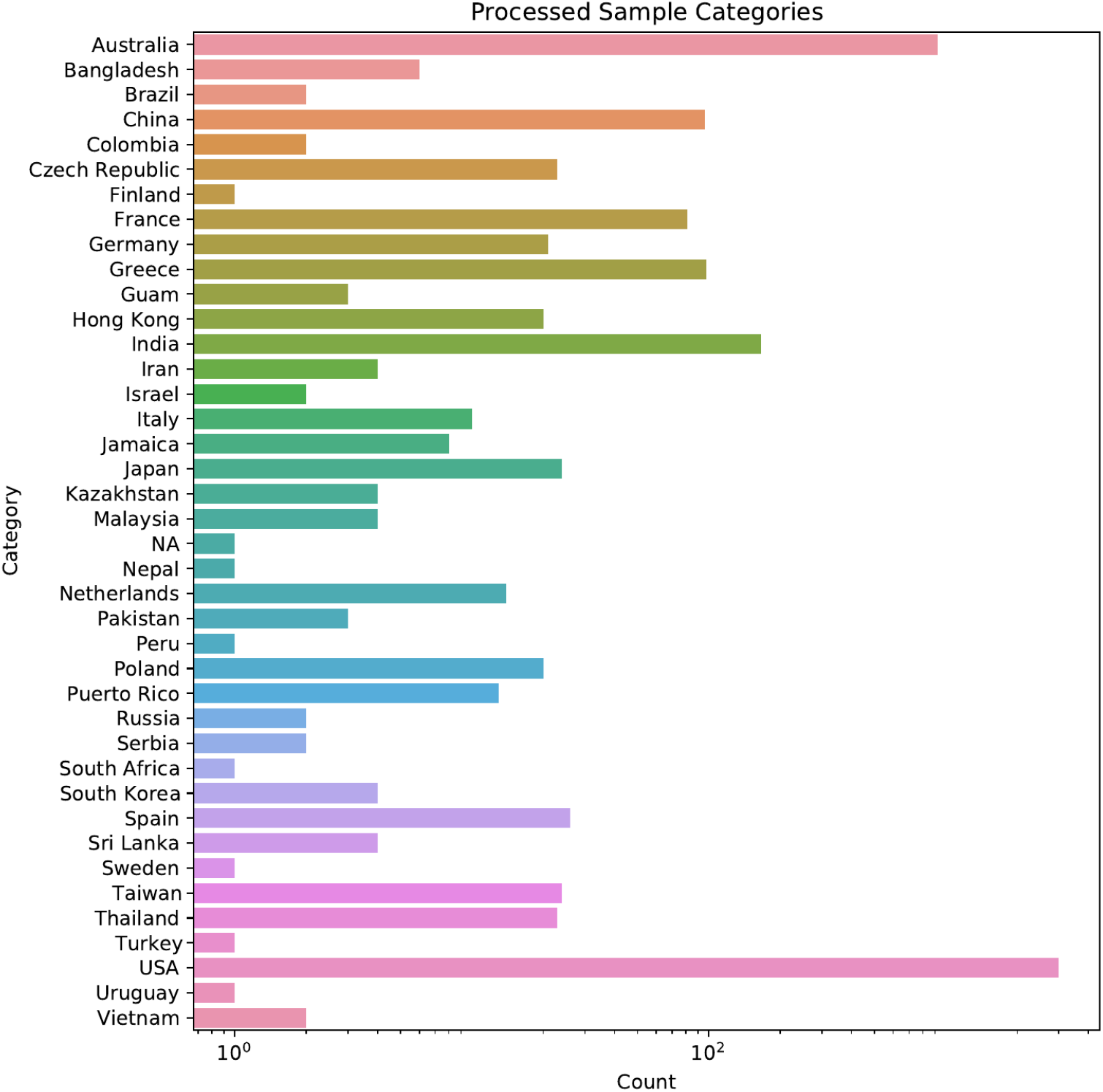
Distribution of categories (country of origin).

We can see from the distribution that nearly 40 countries were represented in the dataset, with the five highest sequence counts from the U.S., Australia, Hong Kong, Germany, and China. Incidentally, as of June 2020, the top five countries in terms of total number of confirmed cases are the U.S., Brazil, Russia, India, and the United Kingdom [3]. While there does not seem to be significant correlation between sequence counts and confirmed cases, which could be the result of sampling bias, it is notable that the U.S. has the highest counts for both metrics.

Multiple sequence alignment resulted in 29903 positions (213 invariant) and 4333 unique sequences. The average pairwise sequence distance was 0.000323, with a standard deviation of 0.000181.

**Figure 6:**
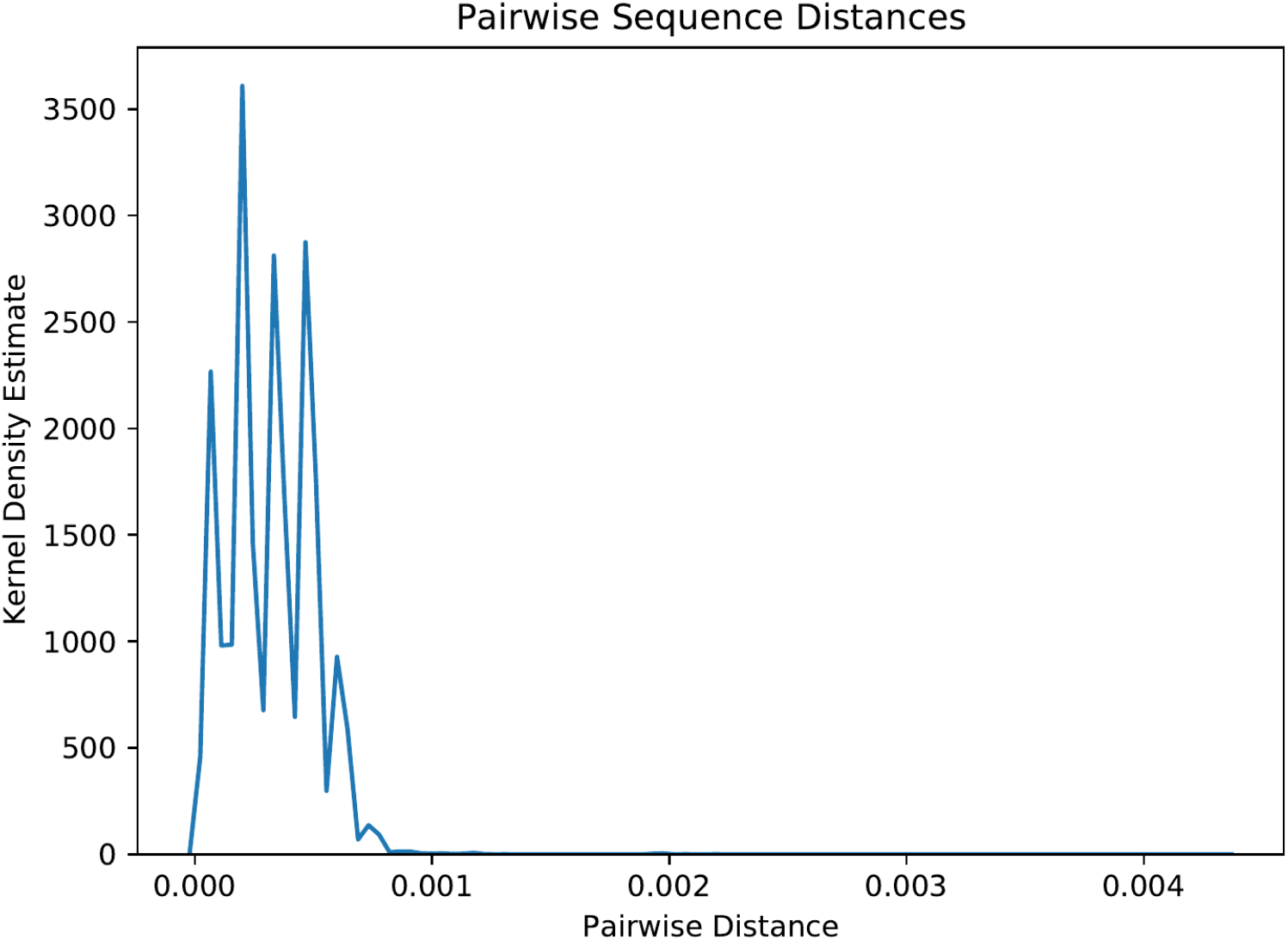
Distribution of pairwise sequence distances.

Pairwise sequence distances indicate how closely related the sequences are, and in comparison to the average pairwise sequence distance from the various coronavirus strains (Section 3.1) of 0.503, the average for the SARS-CoV-2 genomes of 0.000323 is multiple orders of magnitude smaller, indicating that the SARS-CoV-2 genomes are much more closely related to each other, as expected.

Across the multiple sequence alignment positions, the average coverage was 0.983 with a standard deviation of 0.0416, indicating good coverage. The average Shannon entropy as 0.00993 with a standard deviation of 0.0431; these relatively small values indicated that there was not much sequence variability.

**Figure 7:**
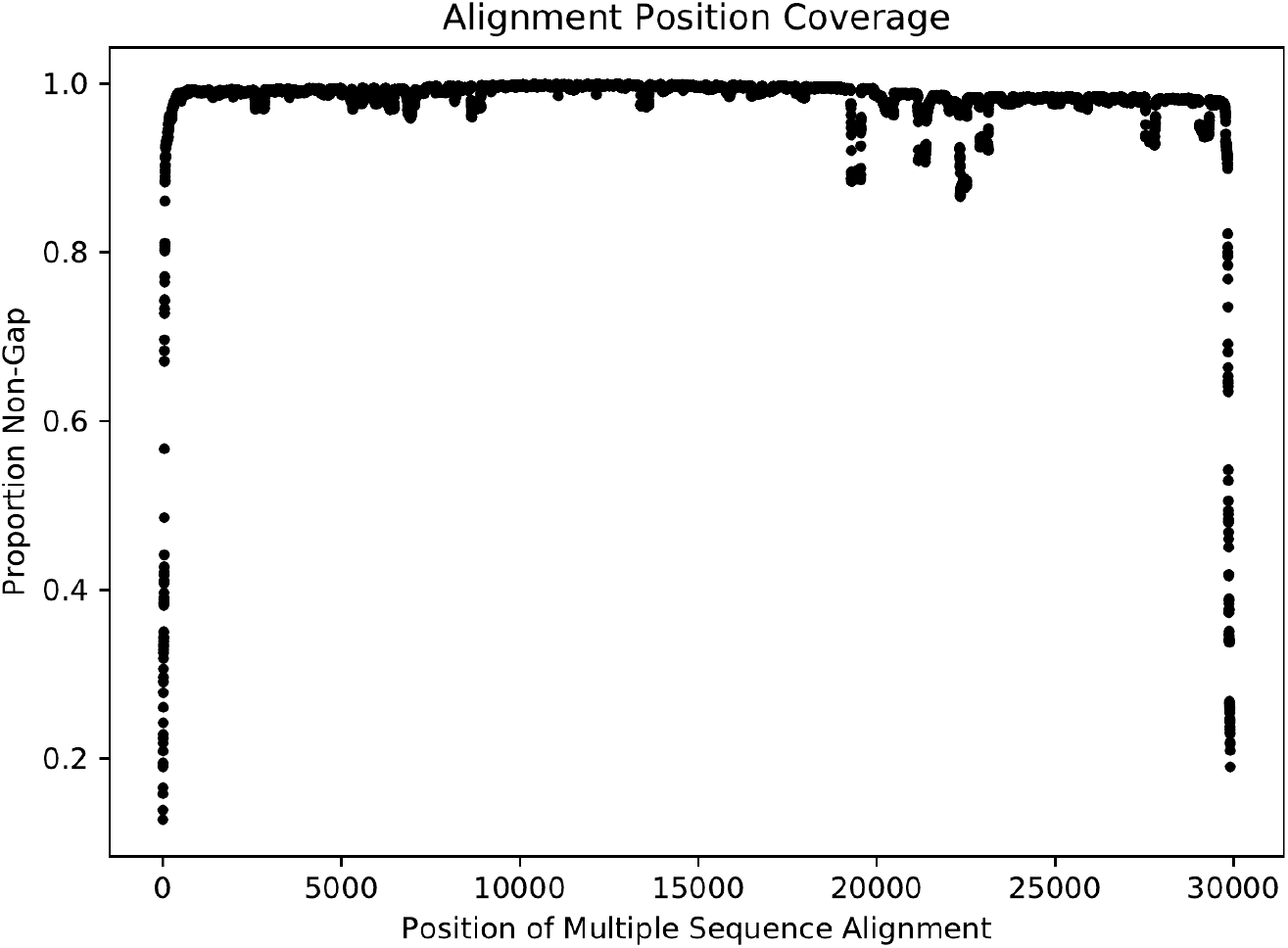
Coverage (proportion of non-gap characters) across MSA positions.

**Figure 8:**
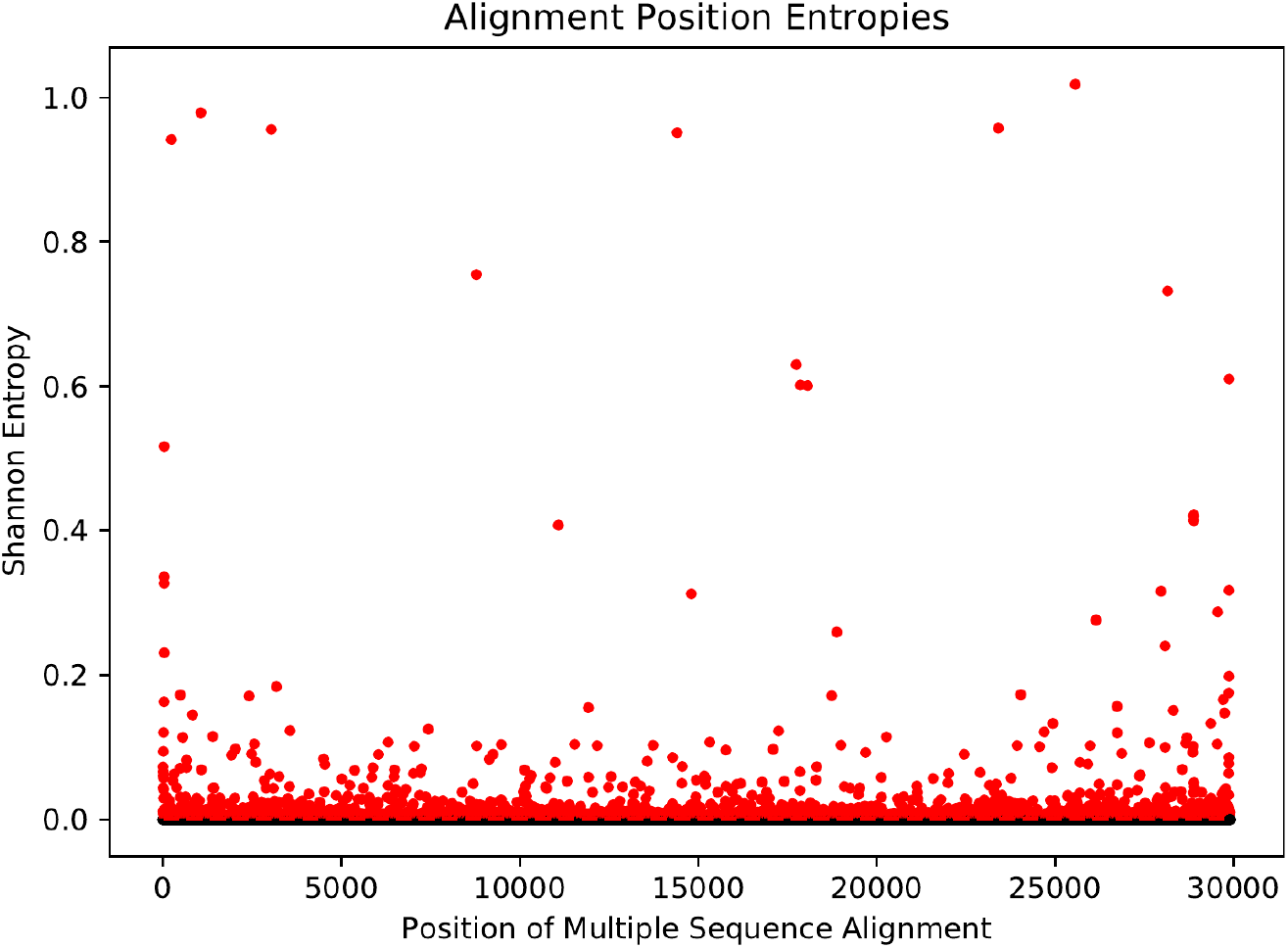
Shannon entropy across MSA positions. Due to the abundance of zero-entropy positions, all non-zero entropies were deemed significant. The significance threshold is shown as a red dashed line, and significant points are shown in red.

The following maximum-likelihood phylogeny was inferred, with an average pairwise phylogenetic distance of 0.000381 and a standard deviation of 0.000208. The maximum pairwise phylogenetic distance (i.e. tree diameter) was 0.00505, which describes the distance between the two least closely related sequences.

**Figure 9:**
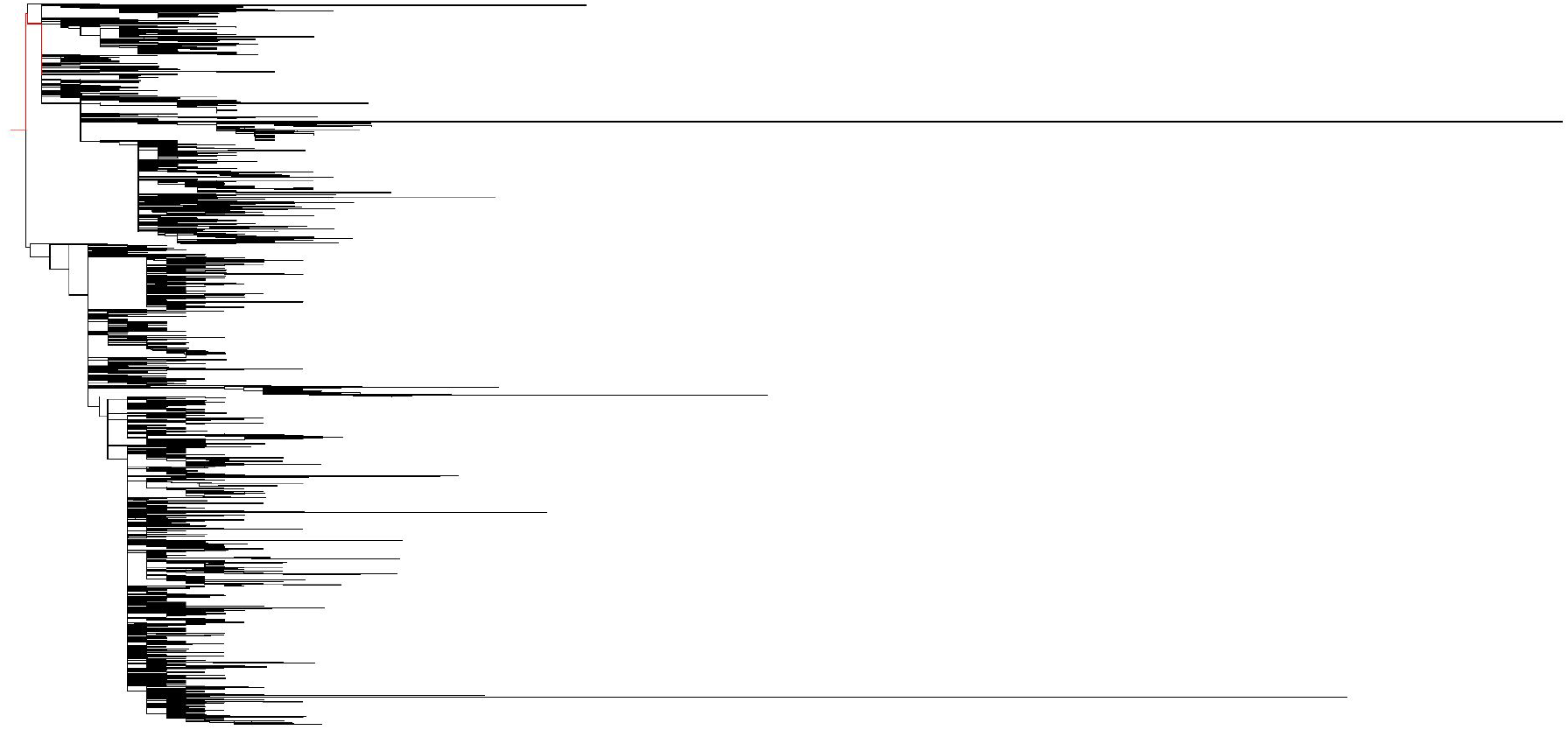
Phylogenetic tree as visualized using IcyTree, path to reference sequence from root highlighted in red.

**Figure 10:**
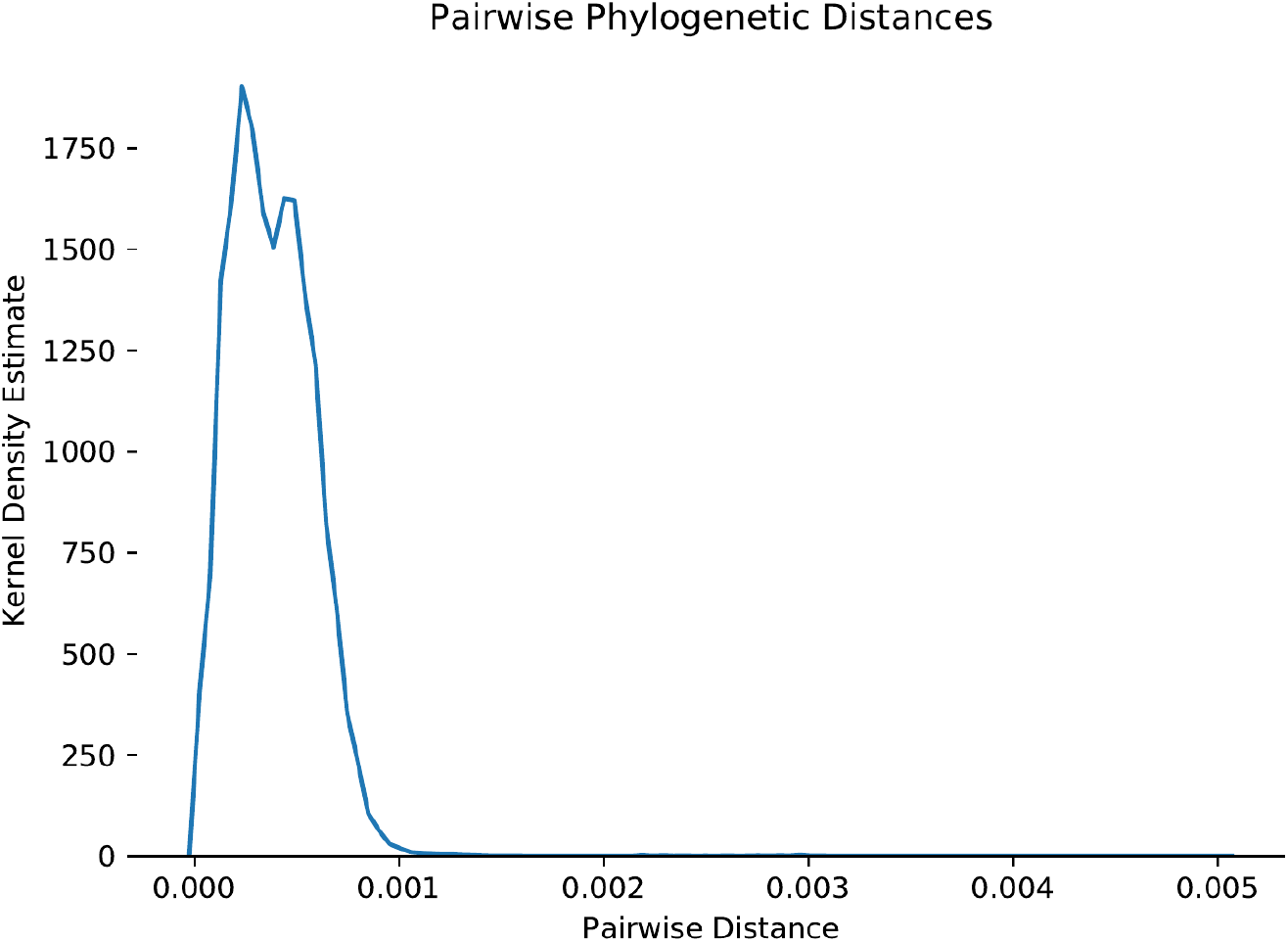
Distribution of pairwise phylogenetic distances.

Given the large number of sequences included, it is evidently difficult to visualize the specific relationships between sequences in the tree. These individual relationships can be visualized by inputting the rooted tree file to iTOL, but the tree is too large to include in this report.

The pairwise phylogenetic distances were also used for transmission clustering using TreeN93 [12], which resulted in 736 clusters and 2535 singletons. The average cluster size was 2.853 with a standard deviation of 2.0284, and cluster sizes ranged from 2-30.

**Figure 11:**
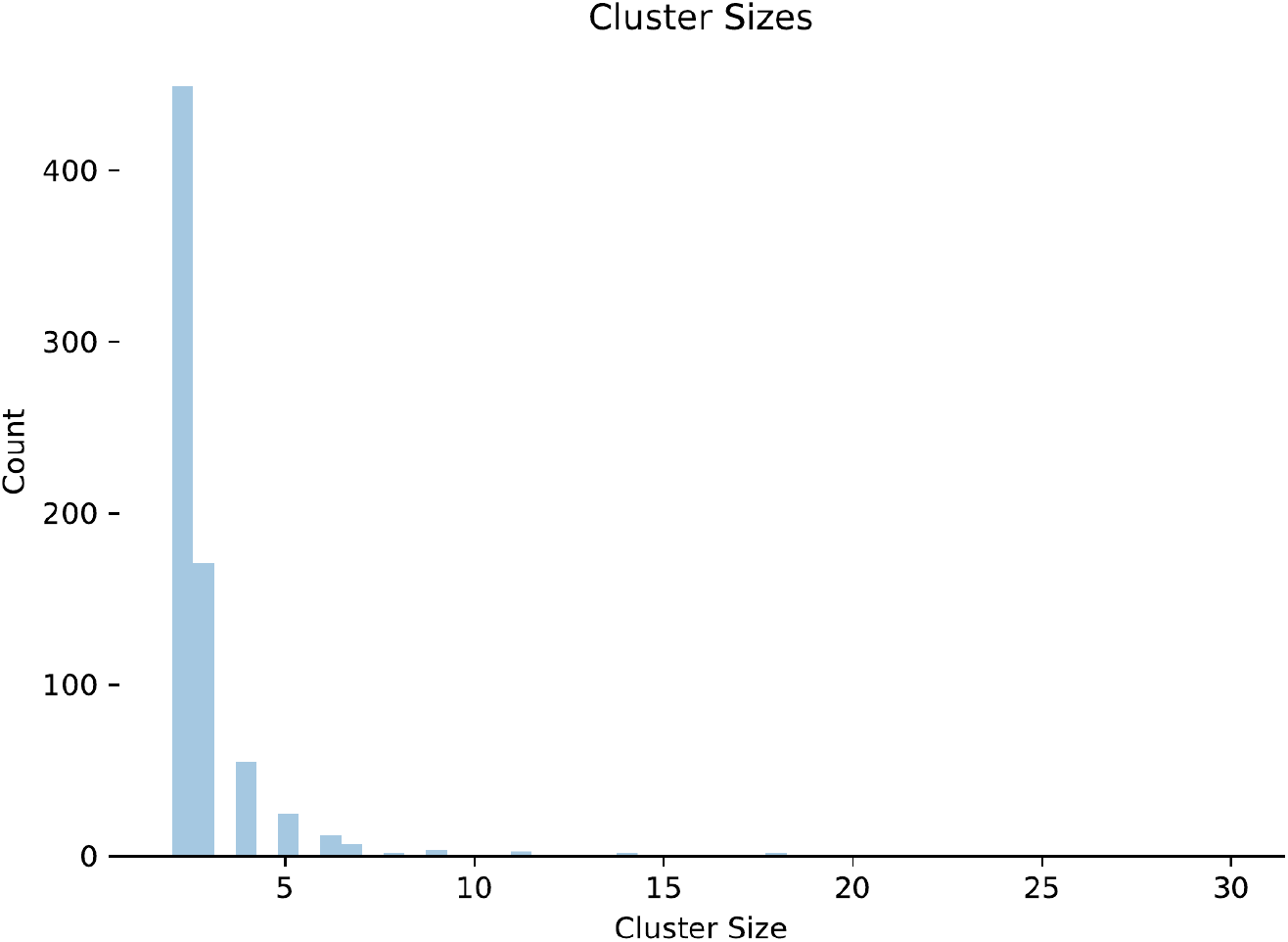
Distribution of cluster sizes.

## 4 Discussion

Some considerations of the limitations of this analysis include the dataset used; given that there are well over 8 million confirmed cases of SARS-CoV-2 in the world (as of June 2020) but the sample only included around 4500 sequences, the sample is evidently a very small subset that may not be representative of current events both in geographical distribution and time. For example, the UK has the fifth most cases in the world [3] but is not listed as one of the categories represented by the sample. As the viral sequence database updates with new sequences or sequences from other sources are pooled together, the analysis may become more representative.

Future directions for this work include additional analysis of the generated statistics; a topic of interest is how the sequences are distributed by date for each country, as this may be indicative of how successful policies in place are in terms of managing the ongoing epidemic. Another topic is exploring the distribution of pairwise distances over time, which may lead to conclusions regarding the mutation rate of the SARS-CoV-2 virus. With regard to ViReport, additional work is being done to analyze potential differences in results based on which tools are selected at each step.

## 5 Data Availability

All genome sequences are available at **NCBI Virus**. The complete ViReport output can be found at https://github.com/mirandajsong/ViReport-SARS-CoV-2.

